# Overlapping Genetic Architecture between Parkinson Disease and Melanoma

**DOI:** 10.1101/740589

**Authors:** Umber Dube, Laura Ibanez, John P Budde, Bruno A Benitez, Albert A Davis, Oscar Harari, Mark M Iles, Matthew H Law, Kevin M Brown, 23andMe Research Team, Melanoma-Meta-analysis Consortium, Carlos Cruchaga

## Abstract

Epidemiologic studies have reported inconsistent results regarding an association between Parkinson disease (PD) and cutaneous melanoma (melanoma). Identifying shared genetic architecture between these diseases can support epidemiologic findings and identify common risk genes and biological pathways. Here we apply polygenic, linkage disequilibrium-informed methods to the largest available case-control, genome-wide association study summary statistic data for melanoma and PD. We identify positive and significant genetic correlation (correlation: 0.17, 95% CI 0.10 to 0.24; *P* = 4.09 × 10^-06^) between melanoma and PD. We further demonstrate melanoma and PD-inferred gene expression to overlap across tissues (correlation: 0.14, 95% CI 0.06 to 0.22; *P* = 7.87 × 10^-04^), and highlight seven genes including *PIEZO1, TRAPPC2L*, and *SOX6* as potential mediators of the genetic correlation between melanoma and PD. These findings demonstrate specific, shared genetic architecture between PD and melanoma that manifests at the level of gene expression.

## Introduction

An association between idiopathic Parkinson disease (PD), neuropathologically characterized by the degeneration of pigmented dopaminergic neurons, and cutaneous melanoma (melanoma), a cancer of pigment-producing melanocytes, was first reported in 1972.^1^ This association was hypothesized to result from the chronic systemic administration of levodopa (L-DOPA) - an intermediate in the dopamine synthesis pathway^2^ – for the treatment of PD^1,3^ as L-DOPA is also a biosynthetic intermediate in the production of melanin.^2^ Since that time, several epidemiologic studies have examined the association between PD and melanoma as well as other cancers.^4–16^ The majority of studies have found that individuals with PD appear to have a lower incidence of most cancers, with the exception of melanoma.^4,5,10,12,13,15,16^ Both prospective and retrospective studies have also found an increased risk of melanoma in PD that appears to be independent of L-DOPA treatment.^4,6,7,9,15^ This increased risk has been observed to extend to family members and be reciprocal in nature with individuals being at greater risk for PD if their relatives have a melanoma diagnosis and vice versa.^6,9^ However, not all studies have identified an association between melanoma and PD in affected individuals^12,17^ or their relatives.^15^ An epidemiologic association between lighter hair color and PD, a potentially shared risk factor with melanoma,^18^ has also been inconsistently reported.^17,19^ Epidemiologic association studies are not without biases. PD is known to have an extended prodromal period and a melanoma diagnosis necessitates longitudinal follow up, both of which increase medical surveillance and thus the chance for spurious epidemiologic associations.^12,20^ In contrast, studies of genetic variants associated with disease or cross-disease risk are not expected to be influenced by usage of medical care, though they may be subject to similar misclassification^21^ and ascertainment biases.

The first investigations of a genetic relationship between melanoma and PD focused on variants in *MC1R*, a gene strongly associated with pigmentation and melanoma risk.^22^ While early reports identified an association between PD and *MC1R* variants^19,23^ other studies failed to replicate these findings.^24–27^ Analyses focused on single variants in other melanoma risk genes have also failed to yield consistent associations with PD.^17,26,28^ Multi-variant analyses have thus far reported a lack of genetic association as well. For example, a melanoma genetic risk score – calculated by aggregating the effect of melanoma genome-wide association study (GWAS)- significant (*P* < 5 × 10^-8^) loci included in the GWAS catalog^29^ as of 2012 – was not significantly associated with PD.^30^ Similarly, no evidence for an association between GWAS-significant melanoma loci and PD is observed in a more recent multi-variant, Mendelian randomization study.^31^ In contrast, genes associated with Mendelian forms of PD have been identified to be somatically mutated in melanoma lesions.^32–34^ There may also exist an enrichment of Mendelian PD gene germline variants in individuals with melanoma,^32^ though this requires replication. Nevertheless, over 90% of individuals with PD do not have mutations in any known Mendelian PD genes.^35^ Thus, variants in Mendelian PD genes are unlikely to fully explain any genetic correlation between melanoma and PD.

The genetic risk architecture underlying complex diseases like PD and melanoma is mediated by many common genetic variants of small effect size, most of which do not demonstrate GWAS-significant associations given current study sample sizes.^36^ Analyses which only include GWAS-significant loci are not expected to fully represent the genetic architecture of these complex diseases and thus may lead to false negative genetic overlap results. Recently, statistical methods that aggregate all associated loci from disease-specific GWAS summary statistic datasets in a linkage disequilibrium (LD)-informed manner have been developed to better model these polygenic architectures.^37^ These aggregated signals can be leveraged to estimate the genetic correlation between different diseases,^37,38^ even at the level of gene expression in specific tissues^39,40^ or across tissues.^41^ Here, we apply these novel methods to GWAS summary statistics derived from the largest currently available studies of melanoma,^22^ PD,^42–44^ and other neurodegenerative diseases^45,46^ to investigate whether specific genetic architecture overlap between melanoma and PD exists.

## Methods

### GWAS Summary Statistics

We obtained the largest available, European genetic ancestry, case-control, GWAS summary statistic data for melanoma (Law2015^22^) and three independent studies of PD (Nalls2014^42^; Chang2017^43^; Nalls2019^44^) as well as two negative control comparator neurodegenerative diseases: Alzheimer disease (Kunkle2019^46^) and frontotemporal dementia (Ferrari2014^45^). The summary statistics for these datasets included p-value, effect allele, number of individuals or studies, and standard error for every genetic variant reported in each study. All individual studies contributing to the GWAS summary statistic datasets used in the current analysis received approval from the pertinent institutional review boards or ethics committees, and all participants gave informed consent. Additional details for each dataset are included below and in the individual study articles.^22,42–46^

#### Melanoma – Law2015

We obtained meta-analysis Melanoma risk summary statistic data from the Melanoma meta-analysis consortium (https://genomel.org/). This data was published in Law et al., Nature Genetics, 2015.^22^ This dataset includes melanoma-association results for 9,469,417 genotyped and imputed variants derived from 12,814 pathologically-confirmed melanoma cases and 23,203 controls of European ancestry.

#### Parkinson disease – Nalls2014

We obtained PD risk summary statistic data from PDGENE (http://www.pdgene.org/). This dataset was published in Nalls et al., Nature Genetics, 2014^42^ and Lill et al, PLoS Genetics 2012.^47^ The summary statistic data we obtained did not include any 23andMe participants and thus the dataset includes PD-association results for 7,799,580 genotyped and imputed variants derived from 9,581 PD cases – mostly diagnosed, but some self-reported – and 33,245 controls of European ancestry. This dataset only included the number of studies, and not the number of individuals, supporting the association results for each variant. Consequently, we only included variants supported by at least 12 of 13 studies in downstream analyses.

#### Parkinson disease – Chang2017

We obtained Parkinson disease (PD) risk summary statistic data from 23andMe, Inc., a personal genetics company (https://research.23andme.com/dataset-access/). This data was published in Chang et al., Nature Genetics, 2017.^43^ This dataset includes PD-association results for 12,896,220 genotyped and imputed variants derived from 6,476 self-reported PD cases and 302,042 controls of European ancestry. This dataset excludes any 23andMe participants included in the Nalls2014 study.

#### Parkinson disease – Nalls2019

We obtained PD risk summary statistic data from the IPDGC (https://pdgenetics.org/). This dataset was provided in the preprint posted by Nalls et al., bioRxiv, 2019.^44^ The summary statistic data we obtained did not include any 23andMe data nor Nalls2014 data and thus includes PD-association results for 17,510,617 genotyped and imputed variants derived from 33,674 PD cases – diagnosed and UKB proxy-cases, that is individuals with a first-degree relative with PD – and 449,056 controls of European ancestry.

#### Alzheimer disease – Kunkle2019

We downloaded stage 1 meta-analysis Alzheimer Disease (AD) risk GWAS summary statistic data from NIAGADS (National Institute on Aging Genetics of Alzheimer Disease Data Storage Site) website: https://www.niagads.org/datasets/ng00075 (#NG00075). This data was generated by the International Genomics of Alzheimer Project and published in Kunkle et al., Nature Genetics, 2019.^46^ The stage 1 meta-analysis dataset includes AD-association results for 11,480,632 genotyped and imputed variants derived from 21,982 AD cases and 41,944 cognitively normal controls of European ancestry.

#### Frontotemporal Dementia – Ferrari2014

We obtained discovery phase Frontotemporal Dementia (FTD) risk GWAS summary statistic data from the International Frontotemporal Dementia Genomics Consortium (IFGC, https://ifgcsite.wordpress.com/data-access/). This data was generated by the IFGC and published in Ferrari et al., Lancet Neurology, 2014.^45^ The discovery phase dataset includes FTD-association results for 6,026,385 variants derived from 2,154 individuals with FTD and 4,308 control of European ancestry.

### Meta-analyzing PD GWAS datasets

We used METAL software^48^ to perform an inverse-variance weighted meta-analysis of the three independent PD GWAS summary statistics. We refer to this meta-analyzed PD dataset in the text, tables, and figures as METAPD (49,731 cases and 784,343 controls).

### Standardization and Filtering of GWAS Summary Statistics

We standardized all summary statistics prior to polygenic analyses. We first confirmed the genome build to be GRCh37, and then annotated variants with dbSNP v151 rs-identifiers and gnomAD^49^ non-Finnish European (NFE) allele frequencies using ANNOVAR software (2018Apr16)^50^. We only included bi-allelic variants with rs-identifiers and in instances where multiple variants shared the same rs-identifiers, we selected the variant that was supported by the largest number of studies and/or the greatest sample size. Finally, we processed and filtered summary statistics using the munge_sumstats.py tool provided with LD Score Regression Software (LDSC)^37^. This processing and filtering removed variants with an effect allele frequency of less than 0.05 in the gnomAD NFE population, variants with strand-ambiguous alleles, variants supported by a low sample size or effective sample (*N*_*eff*_ = 4/(1/*N*_cases_+1/*N*_controls_)) for the meta-analysis,^48^ and variants that were not reported in the HapMap3 study.^51^ The number of variants overlapping across all processed GWAS summary statistic datasets analyzed in the present study are presented in Table 1.

**Table 1.**
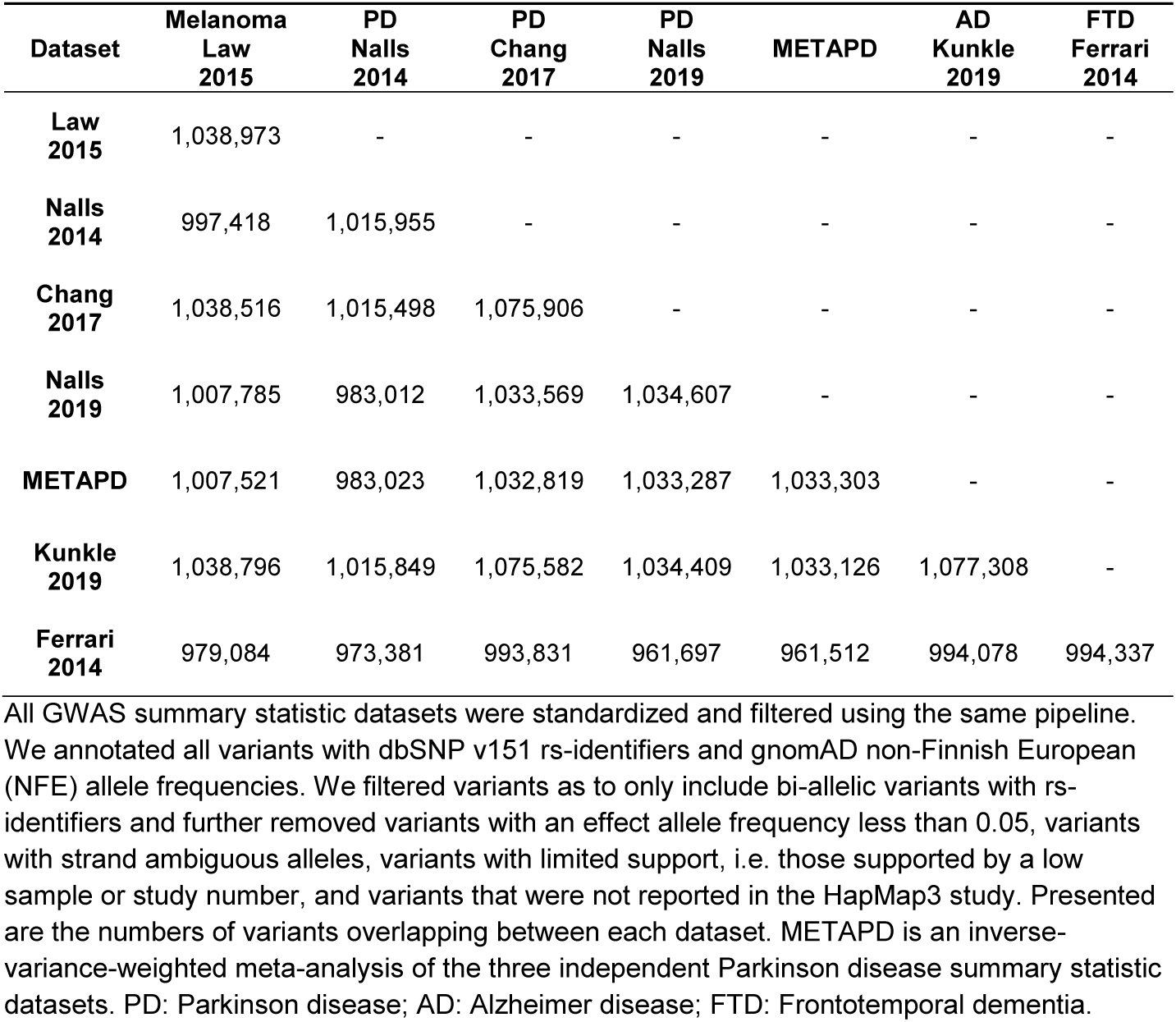
Number of Overlapping Variants in Processed GWAS Summary Statistic Datasets

| **Dataset** | **Melanoma**<br>**Law**<br>**2015** | **PD**<br>**Nalls**<br>**2014** | **PD**<br>**Chang**<br>**2017** | **PD**<br>**Nalls**<br>**2019** | **METAPD** | **AD**<br>**Kunkle**<br>**2019** | **FTD**<br>**Ferrari**<br>**2014** |
| --- | --- | --- | --- | --- | --- | --- | --- |
| **Law**<br>**2015** | 1,038,973 | - | - | - | - | - | - |
| **Nalls**<br>**2014** | 997,418 | 1,015,955 | - | - | - | - | - |
| **Chang**<br>**2017** | 1,038,516 | 1,015,498 | 1,075,906 | - | - | - | - |
| **Nalls**<br>**2019** | 1,007,785 | 983,012 | 1,033,569 | 1,034,607 | - | - | - |
| **METAPD** | 1,007,521 | 983,023 | 1,032,819 | 1,033,287 | 1,033,303 | - | - |
| **Kunkle**<br>**2019** | 1,038,796 | 1,015,849 | 1,075,582 | 1,034,409 | 1,033,126 | 1,077,308 | - |
| **Ferrari**<br>**2014** | 979,084 | 973,381 | 993,831 | 961,697 | 961,512 | 994,078 | 994,337 |

All GWAS summary statistic datasets were standardized and filtered using the same pipeline. We annotated all variants with dbSNP v151 rs-identifiers and gnomAD non-Finnish European (NFE) allele frequencies. We filtered variants as to only include bi-allelic variants with rs-identifiers and further removed variants with an effect allele frequency less than 0.05, variants with strand ambiguous alleles, variants with limited support, i.e. those supported by a low sample or study number, and variants that were not reported in the HapMap3 study. Presented are the numbers of variants overlapping between each dataset. METAPD is an inverse-variance-weighted meta-analysis of the three independent Parkinson disease summary statistic datasets. PD: Parkinson disease; AD: Alzheimer disease; FTD: Frontotemporal dementia.

### Estimating Genetic Overlap by GNOVA

We calculated genetic overlap using GNOVA software^38^. In brief, GNOVA calculates an estimate of genetic covariance between two sets of GWAS summary statistics and further provides an estimate of genetic correlation based on this calculated genetic covariance and the estimated GWAS variant-based heritabilities. As with LD score regression^37^, GNOVA is able to statistically correct for any sample overlap between two different sets of GWAS summary statistics. In addition, GNOVA produces unbounded genetic correlation estimates which may be greater than one for traits which are highly genetically correlated. GNOVA provides greater statistical power and higher estimation accuracy for genetic correlations than LD score regression, especially when the correlations are moderate,^38^ as is expected for melanoma and PD. We ran GNOVA software on the processed GWAS summary statistics using default parameters and the 1000 Genomes^52^ European population-derived reference data provided with the software. Given we test the genetic correlation of melanoma against PD, AD, and FTD we use a Bonferroni corrected significance threshold of *P* < 1.67 × 10^-02^ (0.05 / 3). In the text we present genetic correlations, 95% confidence intervals, and p-values that have been corrected for sample overlap by GNOVA.

### Disease-Inferred Gene Expression Overlap Analyses

We investigated whether the genetic overlap between PD and melanoma was mediated by shared regulation of gene expression. To do this we generated tissue-specific, disease-inferred gene expression profiles from the processed GWAS summary statistics using FUSION/TWAS software.^39^ FUSION/TWAS imputes gene expression using *cis* expression quantitative trait loci (eQTL) data derived from reference panels of paired genotype and tissue-specific gene expression data. As gene expression is imputed based on disease-specific GWAS summary statistics, FUSION/TWAS identifies disease-inferred gene expression profiles with tissue-level resolution. For this study, we used eQTL weights based on the 48 tissue Genotype-Tissue Expression (GTEx)^53^ version 7 (v7) reference panel provided with FUSION/TWAS to generate all disease-inferred gene expression profiles. We tested for overlap or correlation between the disease-inferred gene expression using RHOGE software,^40^ providing the effective sample size^48^ for each dataset. RHOGE provides an estimate of the genetic correlation between two traits that can be attributed to eQTLs as represented by the different trait-inferred gene expression profiles. We exclude the major histocompatibility complex (MHC) region from disease-inferred gene expression overlap analyses due to its complex LD structure.^39,40^ To consider an overlap as significant we used a Bonferroni corrected threshold: *P* < 1.04 × 10^-03^ (0.05 / 48) and present uncorrected p-values and 95% confidence intervals in the text.

### Highlighting Genes Underlying Disease-Inferred Gene Expression Overlap

We used UTMOST software^41^ to generate single-tissue, disease-inferred gene expression, and then aggregated them into a summary metric representing cross-tissue, eGene-disease associations. eGenes are those genes whose expression are influenced by a least one *cis* disease-associated genetic variant.^54^ For this analysis, we generated the single tissue disease-inferred results based on the processed GWAS summary statistics and the 44 tissue GTEx v6 reference panel provided with UTMOST, using default parameters. We similarly generated the cross-tissue summary metric using default parameters. The summary metric generated by UTMOST does not include any indicator of uncertainty. We identified transcriptome-wide significant, cross-tissue, eGene-disease associations using a false discovery rate (FDR) threshold of 0.05, that is five expected false discoveries per 100 reported. We compared PD and melanoma UTMOST summary metric eGene results for the disease-specific GWAS summary statistics to identify eGenes whose expression was similarly regulated by different disease-associated variants.

## Results

### Polygenic Analysis Reveals Specific Genetic Overlap between Melanoma and PD

Prior to cross-disease analyses, we first confirmed that the three independent PD datasets demonstrated positive and significant genetic correlation (genetic correlation range: 0.94 to 1.07, Table 2) using GNOVA software. Following this confirmation and method validation, we proceeded to analyze for potential genetic correlations between melanoma, PD, and the comparator neurodegenerative disease datasets.

**Table 2.** GNOVA Genetic Correlation Results for independent Parkinson Disease Datasets

| Parkinson Disease Dataset | Nalls2014 | Chang2017 | Nalls2019 | METAPD |
| --- | --- | --- | --- | --- |
| **Nalls2014**<br>n<sub>Case</sub> = 9,581<br>n<sub>Control</sub> = 33,245 | - | - | - | - |
| **Chang2017**<br>n<sub>Case</sub> = 6,476<br>n<sub>Control</sub> = 302,042 | 0.95 [0.77, 1.12]<br>(**4.16 × 10<sup>-26</sup>**) | - | - | - |
| **Nalls2019**<br>n<sub>Case</sub> = 33,674<br>n<sub>Control</sub> = 449,056 | 1.07 [0.90, 1.25]<br>(**7.91 × 10<sup>-34</sup>**) | 0.94 [0.80, 1.09]<br>(**1.43 × 10<sup>-36</sup>**) | - | - |
| **METAPD**<br>n<sub>Case</sub> = 49,731<br>n<sub>Control</sub> = 784,343 | 1.00 [0.83, 1.18]<br>(**1.04 × 10<sup>-28</sup>**) | 0.71 [0.56, 0.86]<br>(**8.09 × 10<sup>-21</sup>**) | 1.06 [0.91, 1.21]<br>(**6.10 × 10<sup>-42</sup>**) | - |

We estimated the genetic correlation between the independent Parkinson disease datasets using GNOVA software. All correlation estimates, 95% confidence intervals – presented in square brackets - and p-values - presented in parentheses - are corrected for any potential sample overlap. GNOVA genetic correlation estimates are unbounded and thus may be greater than 1. METAPD is an inverse-variance-weighted meta-analysis of the three independent Parkinson disease summary statistic datasets.

We identified a significant and positive genetic correlation between melanoma and the meta-analyzed PD dataset (genetic correlation: 0.17, 95% CI 0.10 to 0.24; *P* = 4.09 × 10^-06^, Table 3). This result was not driven by any specific PD dataset, but all three independent datasets contributed to the association (*P* < 0.05; genetic correlation range: 0.14 to 0.25, Figure 1 and Table 4). We found no shared genetic architecture between melanoma and Alzheimer disease (genetic correlation: -0.02, 95% CI -0.11 to 0.07; *P* = 0.73, Table 3) nor between melanoma and Frontotemporal dementia (genetic correlation: -0.13, 95% CI -0.37 to 0.12; *P* = 0.32, Table 3).

**Table 3.**
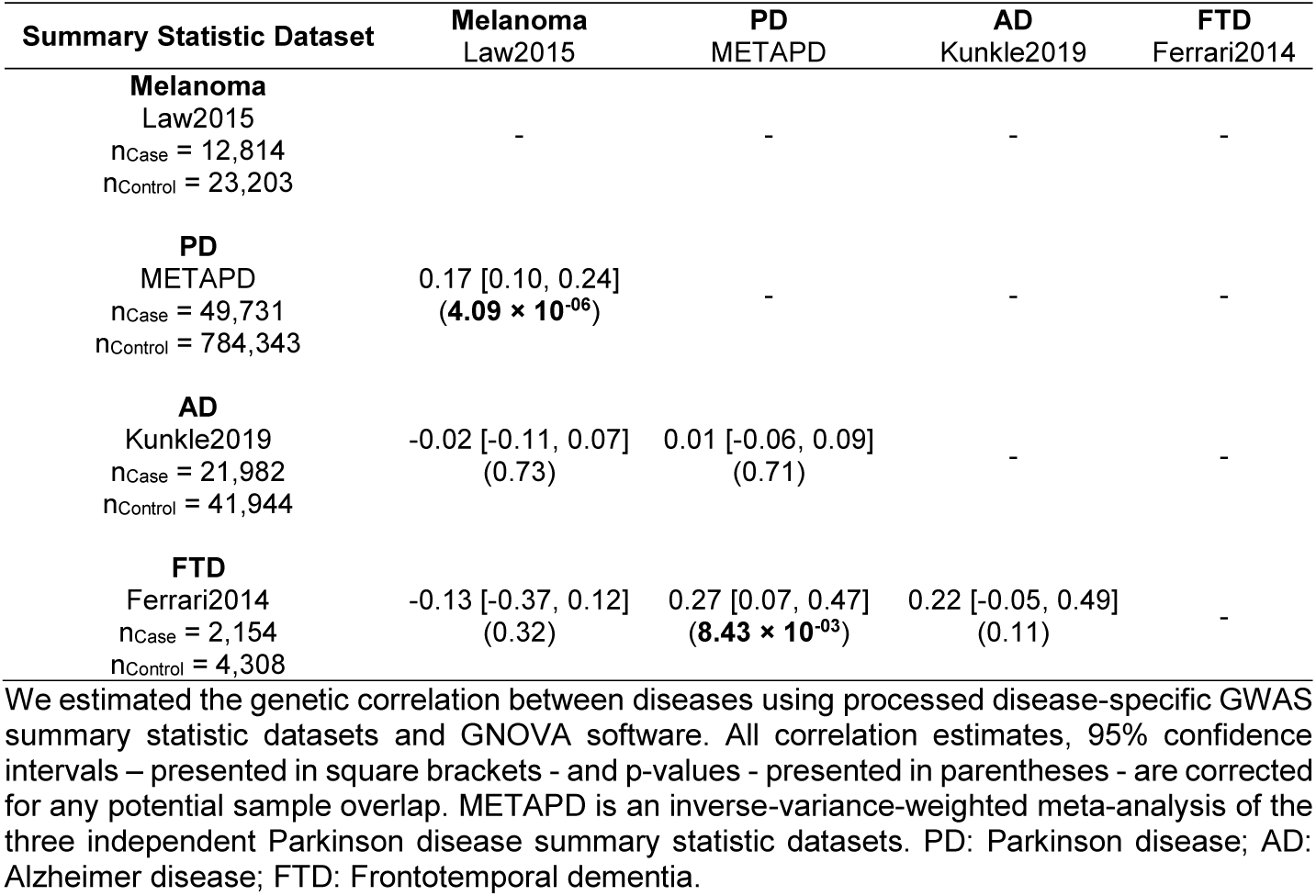
GNOVA Genetic Correlation Results for Meta-analyzed Parkinson Disease, Melanoma, and Comparator Neurodegenerative Diseases GWAS Summary Statistic Datasets

**Table 4.** GNOVA Genetic Correlation Results for Independent Parkinson Disease, Melanoma, and Comparator Neurodegenerative Diseases GWAS Summary Statistic Datasets

| Summary Statistic Dataset | Melanoma<br>Law2015<br>n<sub>Case</sub> = 12,814<br>n<sub>Control</sub> = 23,203 | AD<br>Kunkle2019<br>n<sub>Case</sub> = 21,982<br>n<sub>Control</sub> = 41,944 | FTD<br>Ferrari2014<br>n<sub>Case</sub> = 2,154<br>n<sub>Control</sub> = 4,308 |
| --- | --- | --- | --- |
| **PD**<br>Nalls2014<br>n<sub>Case</sub> = 9,581<br>n<sub>Control</sub> = 33,245 | 0.14 [0.02, 0.25]<br>(**$1.79 \times 10^{-02}$**) | -0.11 [-0.22, 0.00]<br>($4.94 \times 10^{-02}$) | 0.27 [-0.06, 0.60]<br>(0.10) |
| **PD**<br>Chang2017<br>n<sub>Case</sub> = 6,476<br>n<sub>Control</sub> = 302,042 | 0.25 [0.16, 0.33]<br>(**$3.31 \times 10^{-09}$**) | -0.01 [-0.11, 0.09]<br>(0.87) | -0.16 [-0.45, 0.12]<br>(0.26) |
| **PD**<br>Nalls2019<br>n<sub>Case</sub> = 33,674<br>n<sub>Control</sub> = 449,056 | 0.19 [0.10, 0.29]<br>(**$8.28 \times 10^{-05}$**) | 0.05 [-0.04, 0.14]<br>(0.27) | 0.40 [0.14, 0.66]<br>(**$2.78 \times 10^{-03}$**) |

We estimated the genetic correlation between diseases using processed disease-specific GWAS summary statistic datasets and GNOVA software. All correlation estimates, 95% confidence intervals – presented in square brackets – and p-values - presented in parentheses - are corrected for any potential sample overlap. PD: Parkinson disease; AD: Alzheimer disease; FTD: Frontotemporal dementia.

**Figure 1.**
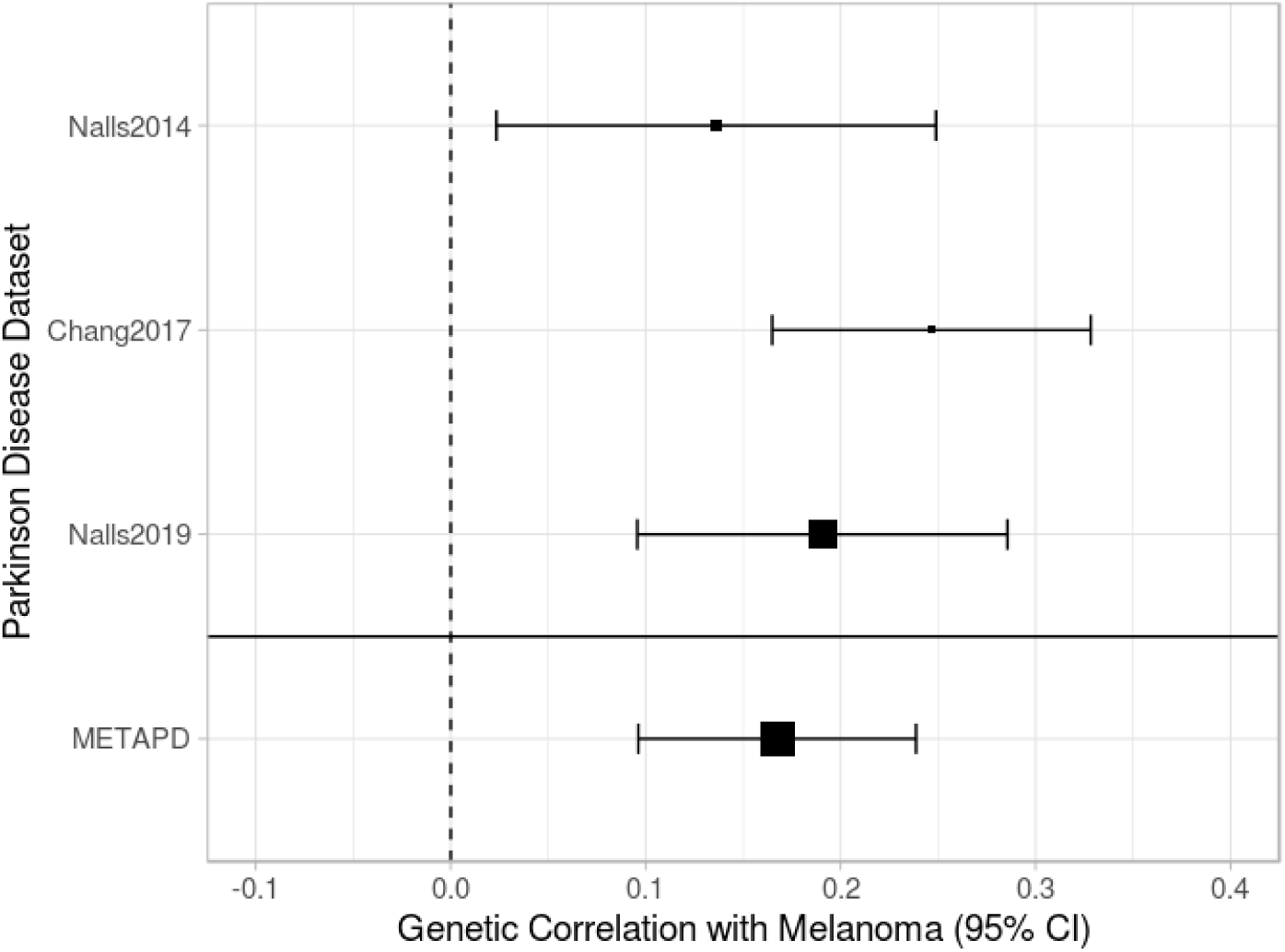
GNOVA Genetic Correlation Results for Parkinson Disease and Melanoma GWAS Summary Statistic Datasets. Forest plot of genetic correlation between melanoma and the individual and meta-analyzed Parkinson disease datasets (Tables 3-4). Box size indicates the effective sample size (*N*_*eff*_ = 4/(1/*N*_cases_+1/*N*_controls_)). The three independent PD datasets are Nalls2014 (Nalls et al., 201442); Chang2017 (Chang et al., 2017^43^); Nalls2019 (Nalls et al., 2019^44^). METAPD is an inverse-variance-weighted meta-analysis of the three independent Parkinson disease summary statistic datasets.

We did not observe any significant correlation between the meta-analyzed PD dataset and AD (Table 3), although one of the individual PD studies showed nominal correlation with AD (Nalls2014: genetic correlation: -0.22, 95% CI -0.22 to 0.00, *P* = 4.94 × 10^-02^; Table 4). We did identify a positive and significant genetic correlation between the meta-analyzed PD dataset and FTD (genetic correlation: 0.27, 95% CI 0.07 to 0.47; *P* = 8.43 × 10^-03^, Table 3), but this appeared to be primarily driven by one of the individual PD studies (Table 4).

Together these results demonstrate a consistent, positive and significant genetic correlation between melanoma and PD but not between melanoma and FTD or AD.

### PD and Melanoma Disease-Inferred Gene Expression Overlaps Across Tissues

To investigate whether melanoma and PD-associated risk variants regulated the expression of the same genes, we generated disease-inferred, tissue-specific gene expression profiles from the processed melanoma and METAPD GWAS summary statistic datasets via FUSION/TWAS software.^39^ We further investigated for overlap between the different disease-inferred gene expression profiles using RHOGE software.^40^

We identified a positive and significant overlap between the PD- and melanoma-inferred gene expression profiles in a joint analysis of the 48 tissues included in the GTEx v7 reference panel provided with the FUSION/TWAS software (disease-inferred gene expression correlation: 0.14, 95% CI 0.06 to 0.22; *P*: 7.87 × 10^-04^). Analyzing the PD- and melanoma-inferred gene expression correlation in each of the reference panel tissues individually, we observed positive overlap in 44 tissues (disease-inferred gene expression correlation median: 0.25, IQR: 0.13, Figure 2 and Table 5), but only a statistically significant overlap in the suprapubic, non-sun-exposed, skin tissue (disease-inferred gene expression correlation: 0.37, 95% CI 0.17 to 0.57; *P*: 7.58 × 10^-04^). Eleven additional tissues demonstrated positive and nominal (Figure 2 and Table 5) the PD- and melanoma-inferred gene expression overlap including spleen (disease-inferred gene expression correlation: 0.40, 95% CI 0.13 to 0.66; *P*: 5.49 × 10^-03^), minor salivary gland (disease-inferred gene expression correlation: 0.45, 95% CI 0.15 to 0.75; *P*: 7.49 × 10^-03^), heart atrial appendage (disease-inferred gene expression correlation: 0.31, 95% CI 0.09 to 0.54; *P*: 8.27 × 10^-03^) brain substantia nigra (disease-inferred gene expression correlation: 0.42, 95% CI 0.14 to 0.71; *P*: 9.02 × 10^-03^), and brain caudate nucleus (disease-inferred gene expression correlation: 0.29, 95% CI 0.01 to 0.58; *P*: 4.89 × 10^-02^).

**Table 5.** Disease-Inferred Gene Expression Profile Overlap between Melanoma and PD in GTEx v7 Reference Panel Tissues

| GTEx v7 Tissue | Number of Samples in Tissue Reference Panel | Melanoma vs. METAPD |  |
| --- | --- | --- | --- |
| | | $\rho_{GE}$ | p-value |
| Adipose Subcutaneous | 385 | 0.30 [0.01, 0.59] | $4.82 \times 10^{-02}$ |
| Adipose Visceral Omentum | 313 | 0.23 [-0.03, 0.49] | $9.39 \times 10^{-02}$ |
| Adrenal Gland | 175 | 0.25 [-0.10, 0.59] | $1.73 \times 10^{-01}$ |
| Artery Aorta | 267 | 0.14 [-0.16, 0.44] | $3.64 \times 10^{-01}$ |
| Artery Coronary | 152 | 0.19 [-0.34, 0.71] | $4.93 \times 10^{-01}$ |
| Artery Tibial | 388 | 0.15 [-0.19, 0.49] | $3.93 \times 10^{-01}$ |
| Brain Amygdala | 88 | 0.25 [-0.10, 0.60] | $1.77 \times 10^{-01}$ |
| Brain Anterior cingulate cortex BA24 | 109 | 0.17 [-0.28, 0.62] | $4.58 \times 10^{-01}$ |
| Brain Caudate basal ganglia | 144 | 0.29 [0.01, 0.58] | $4.89 \times 10^{-02}$ |
| Brain Cerebellar Hemisphere | 125 | 0.18 [-0.18, 0.54] | $3.38 \times 10^{-01}$ |
| Brain Cerebellum | 154 | 0.17 [-0.11, 0.45] | $2.32 \times 10^{-01}$ |
| Brain Cortex | 136 | -0.04 [-0.51, 0.43] | $8.75 \times 10^{-01}$ |
| Brain Frontal Cortex BA9 | 118 | -0.05 [-0.58, 0.49] | $8.67 \times 10^{-01}$ |
| Brain Hippocampus | 111 | 0.41 [0.12, 0.70] | $1.15 \times 10^{-02}$ |
| Brain Hypothalamus | 108 | 0.41 [0.07, 0.75] | $3.09 \times 10^{-02}$ |
| Brain Nucleus accumbens basal ganglia | 130 | 0.34 [-0.04, 0.73] | $9.04 \times 10^{-02}$ |
| Brain Putamen basal ganglia | 111 | 0.30 [-0.04, 0.64] | $9.60 \times 10^{-02}$ |
| Brain Spinal cord cervical c-1 | 83 | 0.26 [-0.56, 1.08] | $5.49 \times 10^{-01}$ |
| Brain Substantia nigra | 80 | 0.42 [0.14, 0.71] | $9.02 \times 10^{-03}$ |
| Breast Mammary Tissue | 251 | 0.24 [-0.09, 0.57] | $1.64 \times 10^{-01}$ |
| Cells EBV-transformed lymphocytes | 117 | 0.09 [-0.39, 0.58] | $7.11 \times 10^{-01}$ |
| Cells Transformed fibroblasts | 300 | 0.29 [0.07, 0.51] | $1.35 \times 10^{-02}$ |
| Colon Sigmoid | 203 | -0.01 [-0.44, 0.42] | $9.60 \times 10^{-01}$ |
| Colon Transverse | 246 | 0.24 [-0.10, 0.57] | $1.70 \times 10^{-01}$ |
| Esophagus Gastroesophageal Junction | 213 | 0.28 [-0.00, 0.56] | $5.88 \times 10^{-02}$ |
| Esophagus Mucosa | 358 | 0.13 [-0.17, 0.43] | $3.92 \times 10^{-01}$ |
| Esophagus Muscularis | 335 | 0.24 [-0.02, 0.51] | $7.36 \times 10^{-02}$ |
| Heart Atrial Appendage | 264 | 0.31 [0.09, 0.54] | $8.27 \times 10^{-03}$ |
| Heart Left Ventricle | 272 | 0.08 [-0.24, 0.41] | $6.22 \times 10^{-01}$ |
| Liver | 153 | 0.25 [-0.07, 0.56] | $1.36 \times 10^{-01}$ |
| Lung | 383 | 0.17 [-0.27, 0.60] | $4.54 \times 10^{-01}$ |
| Minor Salivary Gland | 85 | 0.45 [0.15, 0.75] | $7.49 \times 10^{-03}$ |
| Muscle Skeletal | 491 | 0.17 [-0.07, 0.42] | $1.70 \times 10^{-01}$ |
| Nerve Tibial | 361 | 0.27 [-0.00, 0.53] | $5.61 \times 10^{-02}$ |
| Ovary | 122 | 0.30 [-0.12, 0.71] | $1.79 \times 10^{-01}$ |
| Pancreas | 220 | 0.35 [0.04, 0.66] | $3.15 \times 10^{-02}$ |
| Pituitary | 157 | 0.30 [0.00, 0.59] | $5.54 \times 10^{-02}$ |
| Prostate | 132 | 0.08 [-0.33, 0.49] | $7.10 \times 10^{-01}$ |
| Skin Not Sun Exposed Suprapubic | 335 | 0.37 [0.17, 0.57] | $7.58 \times 10^{-04}$ |
| Skin Sun Exposed Lower leg | 414 | 0.29 [-0.01, 0.58] | $5.96 \times 10^{-02}$ |
| Small Intestine Terminal Ileum | 122 | 0.29 [-0.01, 0.58] | $6.71 \times 10^{-02}$ |
| Spleen | 146 | 0.40 [0.13, 0.66] | $5.49 \times 10^{-03}$ |
| Stomach | 237 | 0.34 [0.04, 0.64] | $3.23 \times 10^{-02}$ |
| Testis | 225 | 0.09 [-0.22, 0.39] | $5.78 \times 10^{-01}$ |
| Thyroid | 399 | 0.26 [-0.02, 0.54] | $7.66 \times 10^{-02}$ |
| Uterus | 101 | 0.30 [-0.02, 0.61] | $8.43 \times 10^{-02}$ |
| Vagina | 106 | -0.11 [-0.93, 0.72] | $8.05 \times 10^{-01}$ |
| Whole Blood | 369 | 0.28 [-0.02, 0.57] | $7.38 \times 10^{-02}$ |

We generated disease-inferred gene expression profiles based on standardized and processed GWAS summary statistics using FUSION/TWAS software and the Genotype-Tissue Expression Project (GTEx) v7 reference panel. We further compared the overlap of these disease-inferred gene expression profiles using RHOGE software. METAPD is an inverse-variance-weighted meta-analysis of the three independent Parkinson disease summary statistic datasets. PD: Parkinson disease; $\rho_{GE}$ : correlation coefficient for inferred transcriptomic overlap; BA: Brodmann Area.

**Figure 2.**
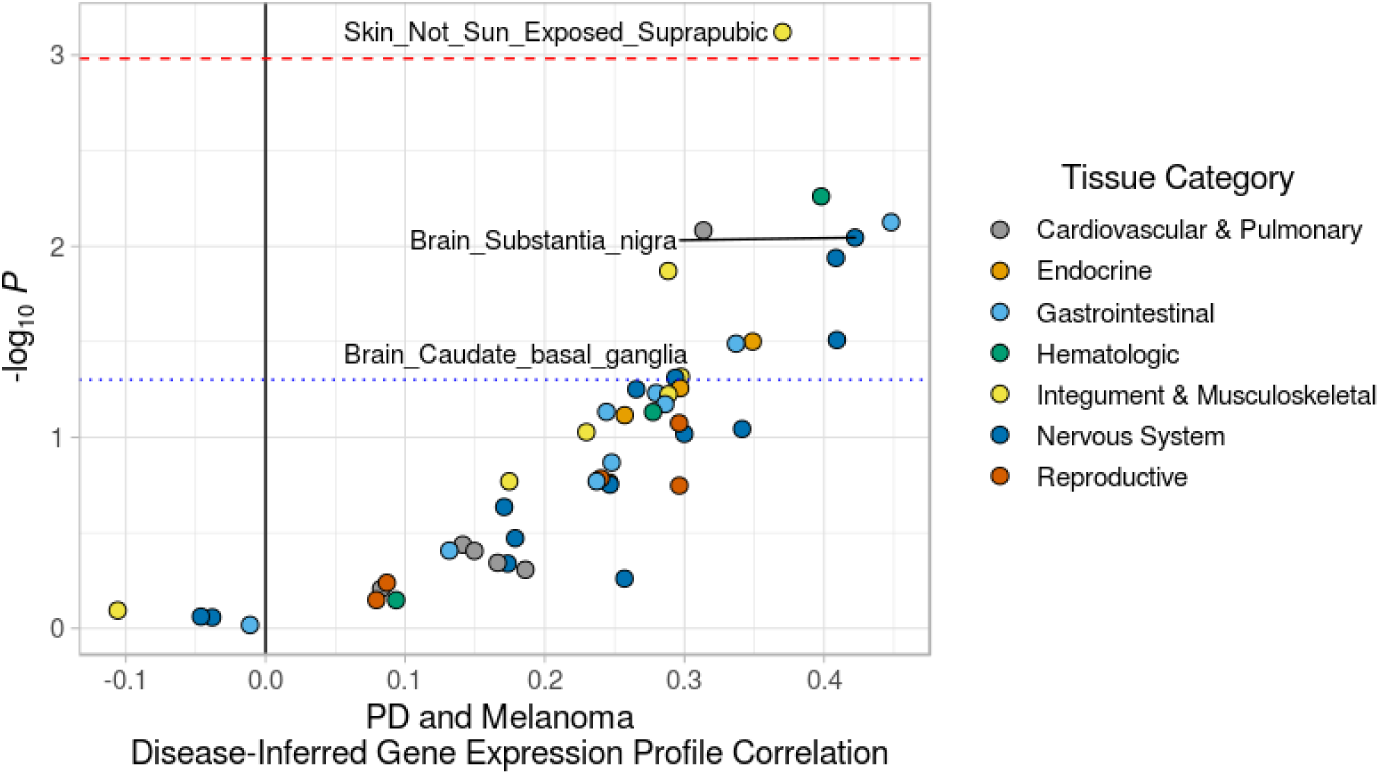
Parkinson Disease (PD) and Melanoma Tissue-specific, Disease-inferred Gene Expression Profile Correlation. PD and Melanoma disease-inferred gene expression profile correlation at the level of 48 specific tissues included in the GTEx v7 reference panel (Table 5). Disease-inferred gene expression profiles were generated from the processed melanoma and METAPD summary statistics using FUSION/TWAS software and correlation between these profiles was estimated using RHOGE software. METAPD is an inverse-variance-weighted meta-analysis of the three independent Parkinson disease summary statistic datasets. The red dashed line demarks the multiple test corrected *P* threshold of 1.04 × 10^-03^ (0.05 / 48) while the blue dotted line demarks the nominal threshold, *P* = 0.05.

To highlight genes whose expression was commonly regulated by PD and melanoma risk variants, we generated cross-tissue, summary metric eGene-disease associations using UTMOST^41^ software. Applying UTMOST to the METAPD GWAS summary statistics, we identified 606 eGenes significantly associated with PD (Supplementary Table 1), including genes in previously reported PD-associated loci^42,47^, such as *MAPT* (*P:* 1.28 × 10^-04^). In the melanoma dataset, we identified 168 significantly associated eGenes (Supplementary Table 2) including those reported in a previous TWAS study^55^, such as *MAFF* (*P:* 1.28 × 10^-12^). Comparing the two sets of cross-tissue summary metric results, we identify seven eGene-disease associations that were passed the FDR threshold for both PD and melanoma: *GPATCH8, MYO9A, PIEZO1, SOX6, TRAPPC2L, ZNF341*, and *ZNF778* (Figure 3 and Table 6). Together, these results suggest that some component of the genetic correlation between melanoma and PD may be mediated by the shared regulation of gene expression across tissues.

**Table 6.** Cross-Tissue eGene-Disease Associations for Melanoma and PD

| Gene | Melanoma | PD |
| --- | --- | --- |
|  | UTMOST Cross-tissue *P* | UTMOST Cross-tissue *P* |
| *GPATCH8* | $8.33 \times 10^{-05}$ | $9.17 \times 10^{-05}$ |
| *MYO9A* | $2.41 \times 10^{-05}$ | $1.01 \times 10^{-03}$ |
| *PIEZO1* | $2.74 \times 10^{-11}$ | $5.65 \times 10^{-05}$ |
| *SOX6* | $1.30 \times 10^{-04}$ | $5.97 \times 10^{-05}$ |
| *TRAPPC2L* | $2.36 \times 10^{-11}$ | $8.47 \times 10^{-05}$ |
| *ZNF341* | $1.67 \times 10^{-04}$ | $1.19 \times 10^{-03}$ |
| *ZNF778* | $2.55 \times 10^{-11}$ | $1.47 \times 10^{-03}$ |

We inferred cross-tissue, eGene-disease associations based on standardized and processed melanoma and METAPD GWAS summary statistics using UTMOST software and the Genotype-Tissue Expression Project (GTEx) v6 reference panel. METAPD is an inverse-variance-weighted meta-analysis of the three independent Parkinson disease (PD) summary statistic datasets.

**Figure 3.**
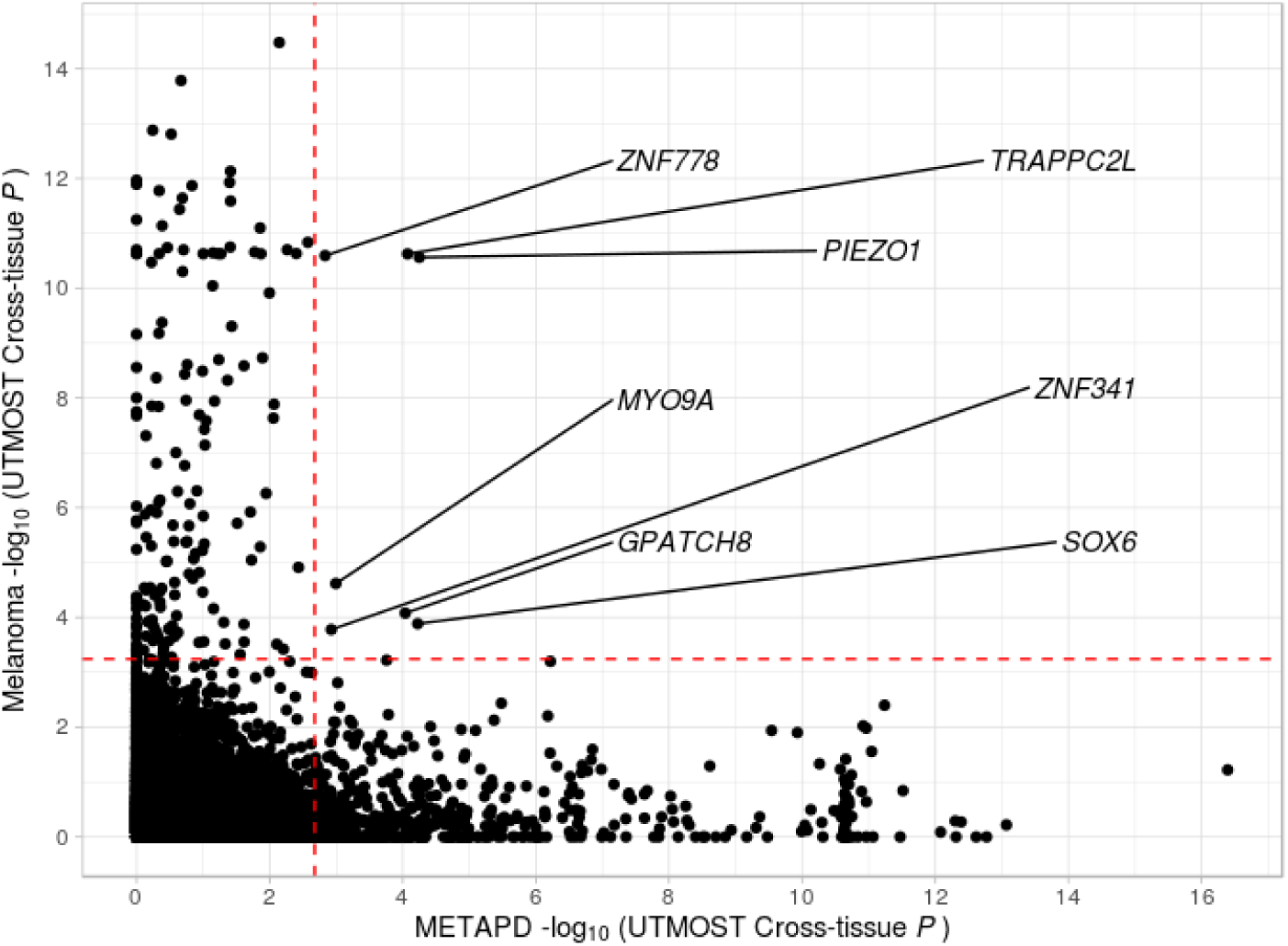
Cross-tissue eGenes Associated with Both Parkinson Disease (PD) and Melanoma. Conjunction plot of the cross-tissue PD and melanoma eGene -log_10_ *P* values. We generated cross-tissue eGene-disease results (Supplementary Tables 1-2) from the processed melanoma and METAPD summary statistics using UTMOST software. METAPD is an inverse-variance-weighted meta-analysis of the three independent Parkinson disease summary statistic datasets. The red dashed lines demark the false discovery rate (FDR) threshold of 0.05. Labels and lines indicate eGenes associated with both PD and melanoma under the FDR threshold.

## Discussion

In this study, we have identified a positive and significant genetic correlation between melanoma and PD by leveraging the largest available GWAS summary statistic datasets and recent advances in polygenic complex trait modeling^37,38^ (Tables 3-4). Our results support the findings of several epidemiologic studies of shared – individual and familial – risk^4–16^ between the two diseases. We also demonstrate no evidence for shared genetic overlap between melanoma and two negative comparison neurodegenerative diseases: AD and FTD (Table 3), suggesting specificity.

Our results of positive genetic correlations between melanoma and PD stand in contrast to negative results from several other genetic studies including single-variant analyses^24–27,30,31^ and multi-variant analyses.^30,31^ Both melanoma and PD are complex diseases with inherently polygenic risk architectures. Consequently, efforts to identify shared genetic architecture at the single-variant level are likely underpowered, especially given the moderate epidemiologic and genetic, correlation between melanoma and PD. This is especially true given the fact that the GWAS results analyzed for such single-variant level investigations are themselves currently underpowered. For example, a power analysis reported in the largest PD GWAS to date (Nalls2019), suggests that an adequately powered PD GWAS would require the inclusion of approximately 99,000 PD cases – more than double their current PD case sample size.^44^ Consequently, our current knowledge regarding the genetic architectures of PD and melanoma is hardly comprehensive and larger GWAS may reveal shared individual risk loci between these diseases in the future. Similarly, previous multi-variant genetic analyses investigating melanoma and PD have focused specifically on GWAS-significant loci and thus can be expected to have missed a substantial proportion of the genetic architecture^36^ underlying these complex diseases. Genetic correlation methods that consider linkage disequilibrium structure and incorporate all associated common variants are better powered to detect genetic overlap, especially given current GWAS sample sizes, as we demonstrate here for melanoma and PD.

The classification and ascertainment of participants was different between the three independent PD datasets included in the present study, but they all demonstrate positive and significant genetic overlap with each other (Table 2). While this overlap does not guarantee specificity of the represented genetic architecture^56^, the fact we observe all three independent PD studies to demonstrate positive and significant genetic overlap with melanoma (Figure 1 and Table 4) bolsters confidence in our results. Importantly, although the PD and melanoma genetic correlation point estimates for the three individual PD studies appear different, their 95% confidence intervals overlap which indicates that the effect size estimates are not significantly different (Figure 1 and Table 4). The genetic overlap between the independent PD datasets supported their meta-analysis, and the genetic correlation between the meta-analyzed PD dataset and melanoma provided the most precise estimate (genetic correlation: 0.17, 95% CI 0.10 to 0.24; *P* = 4.09 × 10^-06^; Figure 1 and Tables 3-4). Further increases in precision can be expected from incorporating additional independent GWAS summary statistic datasets and thus our analyses should be repeated as these become available for both melanoma and PD. Similarly, our FTD genetic correlation results should be interpreted with caution as the current sample size is at least one order of magnitude smaller than the other disease datasets. For example, among the individual PD datasets, we only observe a positive genetic correlation between FTD and Nalls2019. Parkinsonism has been observed in about 20% on individuals with FTD^57,58^, and this result may suggest that individuals with FTD with parkinsonism were included among the UKB-proxy cases in the Nalls2019 dataset. Alternatively, a positive genetic correlation between FTD and the other PD datasets may be observed from the use of a larger FTD GWAS summary statistic dataset. Thus, our analyses should be repeated as larger GWAS summary statistic datasets become available.

We infer disease-associated gene expression profiles^39^ using melanoma and meta-analyzed PD GWAS summary statistics and investigate for their overlap at the level of tissues^40^ and genes^41^ to provide bioinformatically-driven biological context to our melanoma and PD genetic correlation results. We identify significant cross-tissue overlap (disease-inferred gene expression correlation: 0.14, 95% CI 0.06 to 0.22; *P*: 7.87 × 10^-04^) and significant individual tissue overlap in suprapubic non-sun-exposed skin (disease-inferred gene expression correlation: 0.37, 95% CI 0.17 to 0.57; *P*: 7.58 × 10^-04^). We also observe positive, nominal disease-inferred gene expression correlation in peripheral tissues with PD relevance like the heart atrial appendage (disease-inferred gene expression correlation: 0.31, *P* < 0.05, Table 5) - which may reflect the cardiac sympathetic denervation associated with PD^59,60^ - or the minor salivary glands (disease-inferred gene expression correlation: 0.45, *P* < 0.05, Table 5) - which have been reported in some, but not all, studies as containing alpha synuclein aggregates in the context of PD^61,62^. In terms of PD-relevant brain tissues, we observe positive, nominal disease-inferred gene expression correlation in the substantia nigra and basal ganglia caudate nucleus (disease-inferred gene expression correlation: 0.42 and 0.29, respectively; *P* < 0.05, Figure 2 and Table 5). Importantly, the available GTEx v7 inferred gene expression reference model for brain tissues are based on substantially fewer samples than most peripheral tissues, for example the brain substantia nigra reference is derived from 80 donors compared to 335 donors for the suprapubic skin reference (Table 5). Thus, our disease-inferred gene expression risk profile overlap analyses should be repeated as larger reference panels become available.

We identify seven cross-tissue, eGene-disease associations passing the FDR threshold for both melanoma and PD (Figure 3 and Table 6). Importantly, the UTMOST software currently only provides a compatible reference panel based on the GTEx v6 release which is derived from fewer donor samples per tissue compared to GTEx v7 release. In addition, the GTEx v6 reference panel does not include four tissues - brain substantia nigra, brain spinal cervical spinal cord, brain amygdala, and minor salivary gland - which we observed to demonstrate positive disease-inferred gene expression overlap for melanoma and PD (Table 5). Additional eGenes may pass the FDR threshold for both PD and melanoma in analyses based on the larger GTEx v7 reference panel. Thus, our analyses should be repeated when this or other larger reference panels become available for UTMOST. Nevertheless, using the smaller GTEx v6 reference panel we identify seven genes that may be commonly regulated by melanoma and PD-associated variants under the FDR threshold (Figure 3 and Table 6), including *PIEZO1* (Melanoma *P*: 2.74 × 10^-11^; METAPD *P*: 5.65 × 10^-05^); *TRAPPC2L* (Melanoma *P*: 2.36 × 10^-11^; METAPD *P*: 8.47 × 10^-05^); and *SOX6* (Melanoma *P*: 1.30 × 10^-04^; METAPD *P*: 5.97 × 10^-05^).

*PIEZO1* encodes a recently described mechanosensitive cation channel^63^ with several biological functions including human T cell activation^64^ direction of lineage choice in human neural stem cells^65^, and mediating the age-related loss of function of oligodendrocyte progenitor cells^66^. *PIEZO1* is expressed in the neurons of the human substantia nigra^67,68^ and also is ubiquitously expressed in human enteric neurons,^69^ both neuronal types impacted by PD.^70,71^ Interestingly, the expression of *PIEZO2* – *PIEZO1*’s paralog – is regulated by, putatively melanocyte-derived, dopamine signaling in mouse primary sensory neurons^72^ but whether this regulation is relevant for *PIEZO1* is currently unknown.

*TRAPPC2L* is a component of transport protein particle (TRAPP) complexes which function in intracellular vesicle-mediated transport and autophagy.^73–75^ This gene is expressed in human substantia nigra neurons^68^ and a homozygous missense variant in it causes a neurodevelopmental disorder characterized by progressive encephalopathy and episodic rhabdomyolysis.^75^ The intergenic variant rs12921479 - which is an eQTL for *TRAPPC2L* in the brain^53,76^ – was reported to be associated with PD (*P:* 9.31 × 10^-07^) in an autopsy-confirmed cohort of PD^77^, but is only nominally associated with PD in our meta-analyzed PD dataset (*P:* 1.01 × 10^-02^).

*SOX6* is a transcription factor which was recently identified as a determinant of substantia nigra neuron development and maintenance.^78^ Its expression was observed to localize to pigmented and tyrosine hydroxylase positive neurons but not to pigment-negative neurons within the substantia nigra.^78^ In addition, *SOX6* expression was diminished in the substantia nigra of individuals with PD and deletion of *SOX6* in mice was observed to decrease dopamine levels and innervation in the striatum,^78^ a brain region that is also impacted in PD.^79^ In a separate study, a large deletion in *SOX6* was identified in a patient with global developmental delay and progressive parkinsonian symptoms including rest tremor.^80^ Together, these findings suggest a biologically plausible role for *PIEZO1, TRAPPCL2*, and *SOX6* in the genetic correlation between melanoma and PD, but these findings require confirmation and further investigation with future experimental work.

PD and melanoma are clinically heterogenous diseases^81,82^ for which spatiotemporal environmental exposures are relevant^82,83^ and may be necessary, in addition to innate genetic susceptibility, for the development of sporadic disease. Consequently, the moderate genetic correlation we observe should not be interpreted as suggesting that these diseases will always be co-morbid. However, our results of replicable and significant genetic correlation, regardless of the magnitude of effect, do suggest that these two very different diseases share common biological pathways. Thus, even if only a minority of individuals with PD ultimately develop melanoma, understanding the genetic correlation between these disease at the molecular level – for example, if and how the regulation of *PIEZO1, TRAPPC2L*, and *SOX6* and their related biological pathways contribute to PD etiopathogenesis – may provide mechanistic insight that is generalizable to all individuals with PD. Our results support such future research efforts.

## Acknowledgements

We thank Dr. Susan Searles Nielsen for helpful comments on a previous version of this manuscript.

This work was supported by grants from the National Institutes of Health (R01AG044546, P01AG003991, RF1AG053303, R01AG058501, U01AG058922, K01AG046374, K08NS101118 and R01HL119813), the Alzheimer Association (NIRG-11-200110, BAND-14-338165, AARG-16-441560 and BFG-15-362540). This work was supported by access to equipment made possible by the Hope Center for Neurological Disorders and the Departments of Neurology and Psychiatry at Washington University School of Medicine.

We acknowledge the support of all participants, investigators, and researchers from the Melanoma-Meta-analysis Consortium; complete acknowledgements for this meta-analysis can be found in the supplemental data of Law et al., 2015.^22^

We thank the International Genomics of Alzheimer Project (IGAP) for providing summary results data for these analyses. The investigators within IGAP contributed to the design and implementation of IGAP and/or provided data but did not participate in analysis or writing of this report. IGAP was made possible by the generous participation of the control subjects, the patients, and their families. The i–Select chips was funded by the French National Foundation on Alzheimer disease and related disorders. EADI was supported by the LABEX (laboratory of excellence program investment for the future) DISTALZ grant, Inserm, Institut Pasteur de Lille, Université de Lille 2 and the Lille University Hospital. GERAD/PERADES was supported by the Medical Research Council (Grant n° 503480), Alzheimer Research UK (Grant n° 503176), the Wellcome Trust (Grant n° 082604/2/07/Z) and German Federal Ministry of Education and Research (BMBF): Competence Network Dementia (CND) grant n° 01GI0102, 01GI0711, 01GI0420. CHARGE was partly supported by the NIH/NIA grant R01 AG033193 and the NIA AG081220 and AGES contract N01–AG–12100, the NHLBI grant R01 HL105756, the Icelandic Heart Association, and the Erasmus Medical Center and Erasmus University. ADGC was supported by the NIH/NIA grants: U01 AG032984, U24 AG021886, U01 AG016976, and the Alzheimer Association grant ADGC–10–196728.

We acknowledge the PDGENE investigators of the original study^42^ and Drs Lill and Bertram from PDGene^47^ for sharing vbnthe genetics data used for this study.

We would like to thank the research participants and employees of 23andMe for making this work possible.

## Author Contributions

UD conceived the project, designed the study, collected the data, performed the analyses, interpreted the results, and wrote the manuscript. LI, JPB, BAB, AAD, OH, MMI, MHL, and KB contributed to data collection and result interpretation. CC designed the study, collected the data, supervised the analyses, interpreted the results, and wrote the manuscript. All authors read and contributed to the final manuscript.

